# Discovery and characterization of a pancreatic β cell subpopulation expressing an unknown surface epitope through single cell proteomics

**DOI:** 10.1101/2025.09.21.677593

**Authors:** Y. Moursli, D. Faubert, C. Grou, B. Coulombe

## Abstract

The pancreatic β cell surface proteome is essential for supporting cell communication, as it contributes to their secretory identity and maintains β cell responsiveness to any extra- and intracellular variations. Therefore, studies are actively looking to identify specific surface biomarkers that not only can differentiate dysfunctional cell populations, particularly in the context of diabetes, but also can be used to develop therapeutic targets. While β cell heterogeneity has been explored at the single-cell genomic and transcriptomic levels, in this study we aimed to demonstrate that even when a new plasma membrane epitope is identified, characterization of the proteome of cells expressing the same surface protein confirms that different subpopulations do exist. Using single-cell proteomic approach, we analyzed the proteome of MIN6 and αTc1 cells expressing Cd71, Cd99, and P538, which binds to a Fab-phage-538 identified in our previous study. Our findings suggest that the MIN6 (P538^+^) subpopulation may hold a more specialized functional identity compared to the other MIN6 (Cd71^+^), MIN6 (Cd99^+^), and control subpopulations. Moreover, we demonstrate that cell lines derived from the same tissue and expressing the same surface epitope can exhibit heterogeneity in their proteome. Indeed, while the specific expression of 17 protein groups confers to the MIN6 (P538^+^) subpopulation an insulin production and secretion-oriented identity, αTc1 (P538^+^), on the other hand, was characterized by only two proteins with no established involvement in glucagon production or secretion. Also, while the MIN6 (Cd99^+^) subpopulation may be predisposed to the development of insulin deficiency compared to MIN6 (Cd71^+^) and MIN6 (P538^+^), the proteomic profile of the αTc1 (Cd99^+^) subpopulation suggests that this subpopulation may hold an ability to adapt to stressful environmental conditions. Finally, when comparing MIN6 (Cd71^+^) to αTc1 (Cd71^+^), our findings suggest that the MIN6 subpopulation has probably a lesser role in Type 2 Diabetes Mellitus (T2DM) genesis in the context of iron homeostasis disruption than αTc1 (Cd71^+^).

## Introduction

Surface biomarkers have long been considered ideal pharmacological targets (Stutzer I, 2012). However, as demonstrated by several studies, when a molecule targets a surface epitope expressed ubiquitously in multiple cell lines, the ligand’s effect on the cellular response varies (Deng W, 2024). Differential proteomic capital and composition between cells is one of the explanations proposed (Deng W, 2024). In fact, treatments with anti-diabetic molecules are frequently accompanied by several side effects, which can be explained by the variation in the proteomic profile or by differences in signaling pathways and regulation mechanisms between cell subpopulations (Deng W, 2024).

β cells have a rich and complex surface proteome that interacts with various signals such as hormones, ions, metabolites, peptides, and proteins (Stutzer I, 2012). According to transcriptomic studies, the surface proteomic profile can be affected by pathological microenvironments, leading to the appearance of different β cell subpopulations (Dorrell C, et al., 2016). Furthermore, other studies have demonstrated that the manipulation of intracellular protein expression levels can significantly impact and reshape the surface proteome architecture (Zhang Y, 2010). These findings suggest that the proteomic profile of cells identified by surface binders, can vary even between cells of the same lineage (Rugg-Gunn PJ, 2012). In other words, cell diversity at the proteomic level should be addressed when designing new therapeutic molecules, since cells from the same lineage, or from different tissues, may express the same surface epitope and have distinct proteomic profiles.

Cell heterogeneity is in fact a biological phenomenon with several definitions. Initially, cell diversity in the same tissue was observed within cancerous tissues (Tellez-Gabriel M, 2016) (Elsasser WM, 1984). Over the last decade, the study of pancreatic β cell populations diversity, especially within the context of diabetes mellitus (DM), has received a lot of attention among many research groups (Weng C, 2023). β cell diversity exploration was particularly based on single-cell genomic and transcriptomic approaches. For example, Xin Y. et al. through single cell RNA sequencing on β cells isolated from 12 pancreas of healthy human donors, characterized four distinct subpopulations (Dominguez-Gutierrez G, 2019) (Xin Y, 2018). Analysis of signaling pathways of enriched genes in each subpopulation revealed that one cell group with the highest genetic enrichment (433 genes out of a total of 488), was mainly associated with endoplasmic reticulum (ER) stress response and protein folding (Dominguez-Gutierrez G, 2019) (Xin Y, 2018). Xin Y. et al. findings align with other work emphasizing the ER’s crucial role in regulating a wide range of cellular processes involving protein maturation, transcriptional activity, stress responses and apoptosis (Cnop M, 2007). Back and colleagues also found a link between oxidative stress, ER dysfunction and β cell disorders, emphasizing the need to further investigate the proteomic profile changes of these cells (Back SH, 2009). Proteins are in fact comparable to worker ants in a colony, as they continuously contribute to a cell’s specialization, function, phenotype and response to external stimuli like drugs or pathogens (Arias-Hidalgo C., 2022).

In this study, we explored pancreatic β cells (MIN6) diversity by first identifying an unknown surface epitope through an unbiased cell-based biopanning method using Fab-phage display technology (Moursli Y, 2025). The protein P538 was identified through the high binding properties of the Fab-phage-538 to MIN6 cells compared to αTc1 cells (Moursli Y, 2025). To establish a proof of concept that cells expressing the same surface epitope may exhibit a distinct proteomic profile, even when they belong to the same tissue, we examined the proteomic expression of β cells at the single cell level. The advancement of single-cell proteomic approaches has made possible the quantification and the study of the proteome in individual cells, filling the knowledge gap between transcriptomic data and cell’s functional profile (Slavov N, 2022) (Slavov N., 2021). Using a single cell proteomic method inspired by the ScoPE-MS method (Petelski AA, 2021), we compared the proteome of MIN6 and αTc1 cells expressing Cd71, Cd99, and the newly identified epitope, P538. As anticipated, differences in the proteomic profiles between MIN6 and αTc1 expressing the same epitopes were observed, resulting in the characterization of eight distinct MIN6 and αTc1 subpopulations expressing Cd71, Cd99 and P538. Interestingly, our findings suggest that the MIN6 (P538^+^) subpopulation exhibits the most specialized functional identity associated to insulin production and secretion among all the analyzed subgroups.

## Methodology

### 1. Cell culture

#### The murine pancreatic β and *α* cells culture

MIN6, as β cells, were kindly offered by Dr. Jennifer Estall’s lab (Molecular Mechanisms of Diabetes Research Unit, Montreal Clinical Research Institute), and αTC1-clone 9 (αTC1-c9), as α cells, were purchased from ATCC (10801 University Blvd.., Manassas, VA, US, 20110-2209). Cells were cultured in standard 100 mm petri dishes containing Dulbecco’s Modified Eagle Medium (DMEM) (1X), D-Glucose in 500 ml media from GIBCO (Invitrogen Corporation). The composition of the media for MIN6 included the DMEM (4,5 g/l), 15% of inactivated fetal bovine serum (FBS), 1% of antibiotic solution (5 ml of penicillin and streptomycin), L-glutamine, 110 mg/l of sodium pyruvate and 2,5 ul of 2-Mercaptoethanol (2-ME). For α TC1-c9 cells, the media contained DMEM (3g/l), 10% of inactivated FBS, 1% of antibiotic solution (5 ml of penicillin and streptomycin), L-glutamine, 110 mg/l of sodium pyruvate, 4mM of glutamine, 1% of non-essential amino acids (NEAA), 1,5% of 1M HEPES, and 1ml of 10% BSA.

MIN6 and αTC1 cells were kept in an incubator at humidified 37°C and 95 % air conditions, with 5 % CO_2_ for MIN6 and 10% CO_2_ for αTC1. Every 48 to 72 hours, when cells’ confluence was at 80 to 90 %, MIN6 cells were passaged, however, αTC1cells were passaged every 7 to 12 days depending on their confluence. For routine cell culture maintenance and cell sorting, cells were detached using either cell dissociation enzyme free Hanks’-based Buffer from Gibco, or 0.5mM Ethylenediaminetetraacetic acid (EDTA) in sterile 1x PBS.

### 2. Fluorescence-activated cell sorting - (FACS)-AriaIII

#### Preparation of 96-well LoBind plates for cell sorting

the day before cell sorting, wells of the plates receiving single cells and 1.5 ml Eppendorf Protein LoBind tubes for carrier samples were washed with Optima LC-MS/MS-grade water in a laminar flow hood and incubated overnight at 4^0^C. The next day, the water was removed from wells and tubes, and 2 fmol/μl alcohol dehydrogenase (ADH) was added, and the plates and tubes were incubated overnight at 4^0^C on a shaker. On the day of cell sorting, the ADH solution was removed from each well and tube, and a fresh 1.5 μl of ADH at 2 fmol/μl was added along with 1.5 μl of Optima LC-MS/MS-grade water. The plate was then sealed with a self-adhesive film and kept at 4^0^C until cell sorting.

#### FACS-Aria III cell sorting

MIN6 and αTc1 cells were washed with 5 ml of 1xPBS and detached using 5 ml of 1 x PBS and 0.5 mM of EDTA. The number of cells of each sample varied between 1×10^6^ and 1.5×10^6^ depending on the condition. Cells were sorted based on the expression of Cd71, Cd99, and the protein P538 which binds to a newly identified Fab-phage 538. Thus, MIN6 and αTc1 cells were incubated between 40 to 45 min with primary antibodies anti-Cd71 (Abcam, Cat.ab84036, 1:100), anti-Cd99 (Invitrogen, PA5-102556, 1:200) and anti-M13-HRP (Santa cruz biothechnology, Sc-53004, 1:1000). Cells were then washed with 1xPBS and incubated for 45 min with secondary antibodies Alexa Fluor 647-anti rabbit IgG (Invitrogen, A31573, 1:500) for Cd71 and Cd99 and Alexa Fluor647-anti mouse IgG (Invitrogen, A21235, 1:500) for the phage-Fab_538. Cells were then washed three times with 1 x PBS, resuspended in 300 μl of 1x PBS and kept at 4^0^ C until sorting.

### 3. Cell lysis with mPOP method

The plates and 1.5 ml Eppendorf tubes containing the sorted cells were incubated at −70^0^ C for 1h. In some situations and for logistical reasons, some plates were kept 7 days at −70^0^ C (Petelski AA, 2021) (Petelski AA., 2022). Following this incubation, plates and tubes were heated to 90^0^C using a PCR thermocycler for 10 min with the lid temperature set at 105^0^C then cooled to 12^0^ C (Petelski AA, 2021) (Petelski AA., 2022). Tubes and plates were subsequently sonicated in an ultrasonic bath (SWEEP function) for 5 minutes at room temperature.

### 4. Protein digestion into peptides

Proteins of single cell samples were digested into peptides by adding 1,5 μl of a mixture composed of 1.04 ng/μl trypsin (#V5111, Trypsin modified sequencing grade 100ug, Promega), 0.042 Unit/μl Benzonase Nuclease (#70664-3 10KU from Millipore), and 80 mM Triethylammonium bicarbonate (TEAB). For the carrier samples, 2 μl was added to the tubes from the same mixture but with different concentrations of the trypsin (4.1 ng/ μl) and the Benzonase Nuclease (0.1 Unit/μl). The plates and tubes were then mixed using a vortex for 5 s and centrifuged at 300 rpm. The samples were subsequently incubated at 37°C for 3 h (Petelski AA, 2021) (Petelski AA., 2022).

### 5. TMT labeling

Two TMT plex kits were used, the TMT-11 plex kit (ThermoFisher #UH288615) and the TMT-10 plex kit (ThermoFisher #YG370247). Each TMT-tag reagent had a final concentration of 85 mM (28.85 μg/μl) after solubilization with the acetonitrile (ACN). Each TMT label was then diluted to 1:3 in ACN and 3.3 μl of this dilution was added to the carrier samples. For the single cell samples, each TMT label was diluted to 1:5, and 1 μl was added to the samples. The samples were then incubated at room temperature for 1 h. Then, 1.65 μl of 0.5% hydroxylamine (HA) (HA_50% wt/vol Sigma Aldrich Cat#: 467804-50mL) was added to the carrier samples for a final concentration of 0.071%, and 1 μl of 0.213% HA to the single cell samples, followed by 30 min incubation at room temperature. At the end of the 30 min, the carrier samples were cleaned using Ziptip micropipette tips (C18 resin). From the cleaned final solution, a volume equivalent to 250 cells was added to the single cell samples already marked with TMT. All samples were pooled using the same pipette tip, transferring the volume from each well to the next. The wells were then washed with 5 μl ACN and the samples were dried.

### 6. LC-MS/MS analysis

Pooled TMT-labeled tryptic digests were reconstituted under agitation for 20 min in 8.4 µL of 1.8% ACN and 0.3% formic acid and loaded on a 75 μm i.d. × 250 mm PicoChip C18 column (New Objective) installed on the Vanquish Neo LC system (Thermo Scientific). The buffers used for chromatography were 0.1% formic acid (buffer A), 85% acetonitrile, and 0.1% formic acid (buffer B). Peptides were eluted with a three-slope gradient at a flow rate of 300 nL/min. Solvent B was first increased from 3 to 38% over 67 min, followed by 38 to 60% over 10 min and finally from 60 to 88% over 2 min. The LC system was coupled to an Orbitrap Fusion mass spectrometer (Thermo Scientific) through a PicoChip source (New Objective). Nanospray and S-lens voltages were set to 1.9-2.5 kV and 60 V, respectively. The capillary temperature was set to 250 °C. Full scan MS survey spectra (m/z 360-1550) were acquired in the Orbitrap with a resolution of 60,000 and a target value at 8.8e5. The most intense peptide ions of charge states from 2 to 4 were fragmented by HCD at a normalized collision energy of 30 %. An isolation width of 1.2 m/z was used, and analysis were performed in the Orbitrap with a target value of 1e5 and a maximum injection time of 150 ms. The MS/MS duty cycle was set to 3 s, and target ions selected for fragmentation were dynamically excluded for 6 s.

### 7. Quantitative data processing and cell subpopulations proteomic profiling

The MaxQuant software package (v2.6.5.0) was used to analyze peptide data generated by mass spectrometry. This software was designed to allow quantitative analyses of TMT-labeled peptides based on protein sequences using the entire mouse SwissProt database (Cox J, 2008). All followed MaxQuant parameters steps are provided in table 1 of the supplementary materials.

The R statistical and graphical programming language (version R-4.5.0 for Windows, 64 bits, 86 Mo) was used to analyse and visualize proteomic data generated with MaxQuant. The package Seurat (v5.2.1) was used, particularly codes that are already applied in single cell RNA sequencing data analysis but adapted here to integrate the Reporter Intensity Corrected (RIC). All samples were imported into Seurat and merged. Seurat’s LogNormalize data normalization method was used to reduce batch effects between different injection pools. The expression level of all proteins in the cells was scaled and centered along each protein, and linear dimensionality reduction performed by principal component analysis (PCA). Clusters were identified using Louvain’s algorithm and differential expression analysis on the resulting clusters was conducted using the Wilcoxon rank-sum test with p-values adjusted for multiple testing using the Bonferroni correction.

## Results and discussion

### Cd71 and Cd99 are surface markers expressed by both MIN6 and αTc1

Before conducting single cell proteomic analysis, we first examined the expression of Cd99 and Cd71 epitopes in MIN6 and αTc1 cells. CD99 is a membrane protein encoded by the MIC2 gene and is expressed in some hematopoietic and cancerous cells, like in Ewing’s sarcoma (Manara MC, 2018). CD99 expression in human Langerhans islets has essentially been reported in β and α cells (Martens GA, 2018). In our data, Cd99 expression was detected in both MIN6 and αTc1 with variable intensity, indicating two subpopulations within each cell line. A subpopulation with high Cd99 expression represented 11% and 5% of the total sorted MIN6 and αTc1 cells, respectively. A second subpopulation with lower expression levels represented 71% of MIN6 and 79% of αTc1 sorted cells (figures 2 a, and 2 b). These results are consistent with other findings by Martens et al., which suggest that variations in Cd99 expression level can lead to distinct subpopulations (Martens GA, 2018).

Several mammalian cell lines express the transferrin membrane receptor CD71 which is a type II membrane glycoprotein (Dong HY, 2011). In effort to identify specific cell surface biomarker to distinguish pancreatic β cells from the other Langerhans islet cells, some studies have described the Cd71 as a distinctive biomarker of β cells compared to α cells (Berthault C, 2020). However, our findings conflict with these studies as they indicate that Cd71 was detected in 80% of αTc1 cells as well as in 95% of sorted MIN6 cells (figure 2b). Since the aim of this project was to analyze the proteomic profile of cells expressing the same epitopes, we then proceeded with single cell proteomics profiling of both MIN6 and αTc1 cells based on the confirmed expression of Cd71 and Cd99 proteins (figure 3), as well as the expression of a third surface protein, P538. The latter was recognized by the Fab-phage-538 which was isolated by a cell biopanning method using a Fab-phage display library (Moursli Y, 2025).

### Single cell proteomic analysis allowed the characterization of eight distinct MIN6 and αTc1 subpopulations

The development of a single cell proteomics technique required several trial-and-error tests during the process of adjusting mass spectrometry instrument parameters. We compared the performance of the Orbitrap-Fusion (OF) (Thermo Scientific) and the Q-Exactive (QE) (Thermo Scientific) by evaluating their ability to detect the maximum number of protein groups and to generate the lowest contaminant intensity, which is usually responsible for the background noise. The analysis using OF (Thermo Scientific) yielded the results presented in this paper.

As part of this study, we adapted some components of the SCoPe2-MS approach (Petelski AA, 2021) to our reality to analyze the intracellular proteome of 144 MIN6 and αTc1 cells obtained from 17 experiments (pool injections). The 144 cells correspond to 72 MIN6 and 72 αTc1 cells subdivided as follow: 18 cells from each cell line were sorted based on Cd71, Cd99, P538 protein binding to Fab-phage-538, and the control group represented unmarked cells (table 2 in the supplementary materials). For each unique cell pool, a carrier consisting of 250 cells was used (table in the supplementary materials). In contrast, to the Slavov lab method, we used carrier cells from each cell line and targeted epitope. Using the DDA method, we identified and analyzed 1122 protein groups in 72 MIN6 and αTc1cells (controls and P538), 1101 in 36 MIN6 and αTc1 (Cd99^+^) cells, and 803 in 36 MIN6 and αTc1 (Cd71^+^) cells.

Before analyzing the protein groups detected in each cell, the quality of the preparations and processing of each cell was appreciated. To determine whether the peptide intensities in each TMT channel, representing a single cell, were consistent with the anticipated level, the number of ions reported was measured (Petelski AA, 2021). The total intensities of the reporter ion (RI) of all peptides of a properly prepared cell must be proportional to that of the RI of the peptides of the carrier consisting of 250 cells (Petelski AA, 2021). In our experimental model, this ratio was equal to 1:250, meaning that a valid cell preparation should have an expected relative RI equal to or greater than 0.004 (Petelski AA, 2021). The analysis of the reporter intensity (RI) of each identified peptide in each TMT channel was carried out, after exclusion of contaminants peptides (Petelski AA, 2021) and all 144 cells had a rRI value greater than 0,004 (figure 1 in supplementary materials).

**Figure 1:**
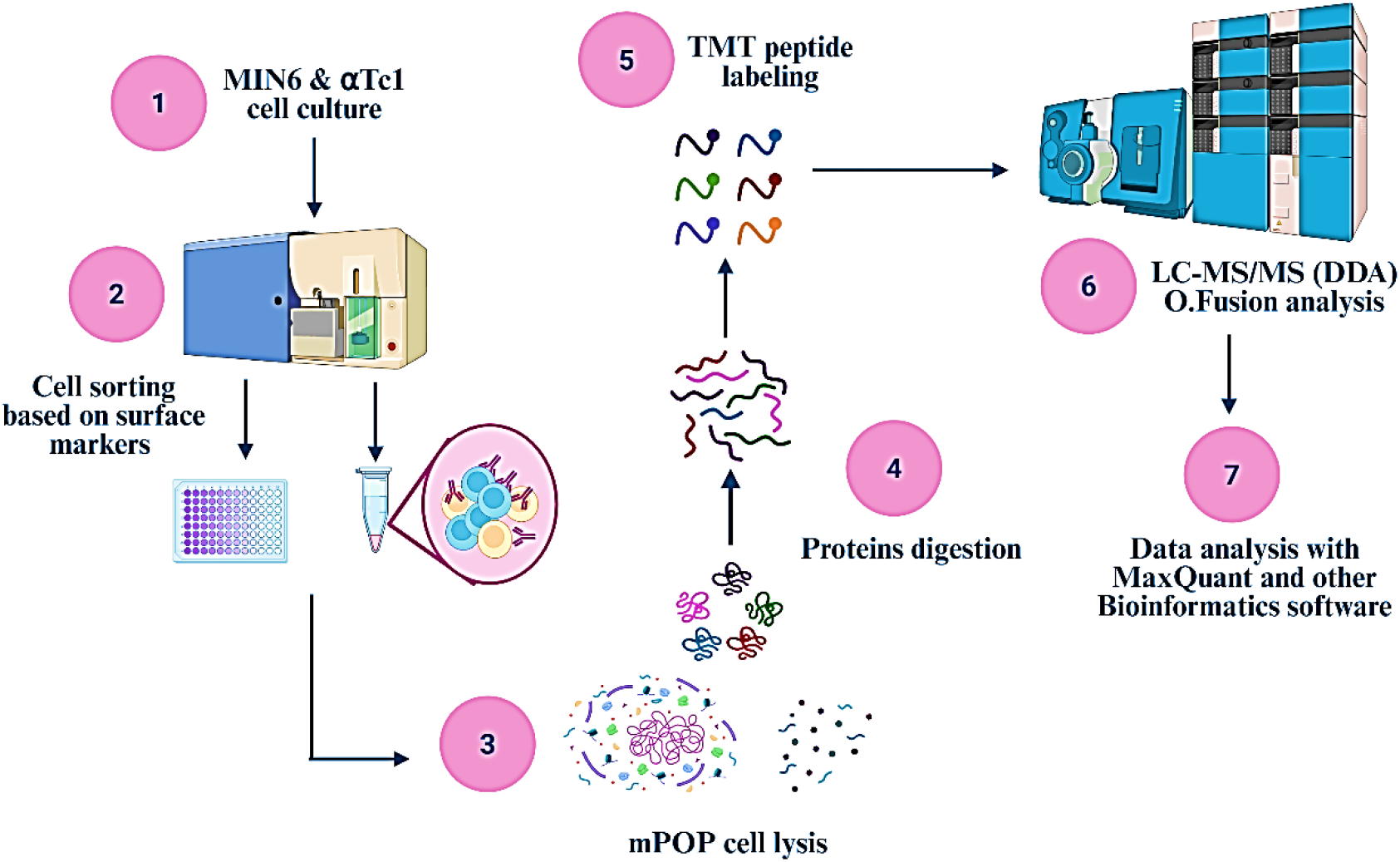
The diagram summarizes the steps involved in developing a single-cell proteomics method inspired by the ScoPe-MS approach (Petelski AA, 2021) (Specht H, 2018), which utilizes cell surface markers and TMT-labeled peptides.

**Figure 2:**
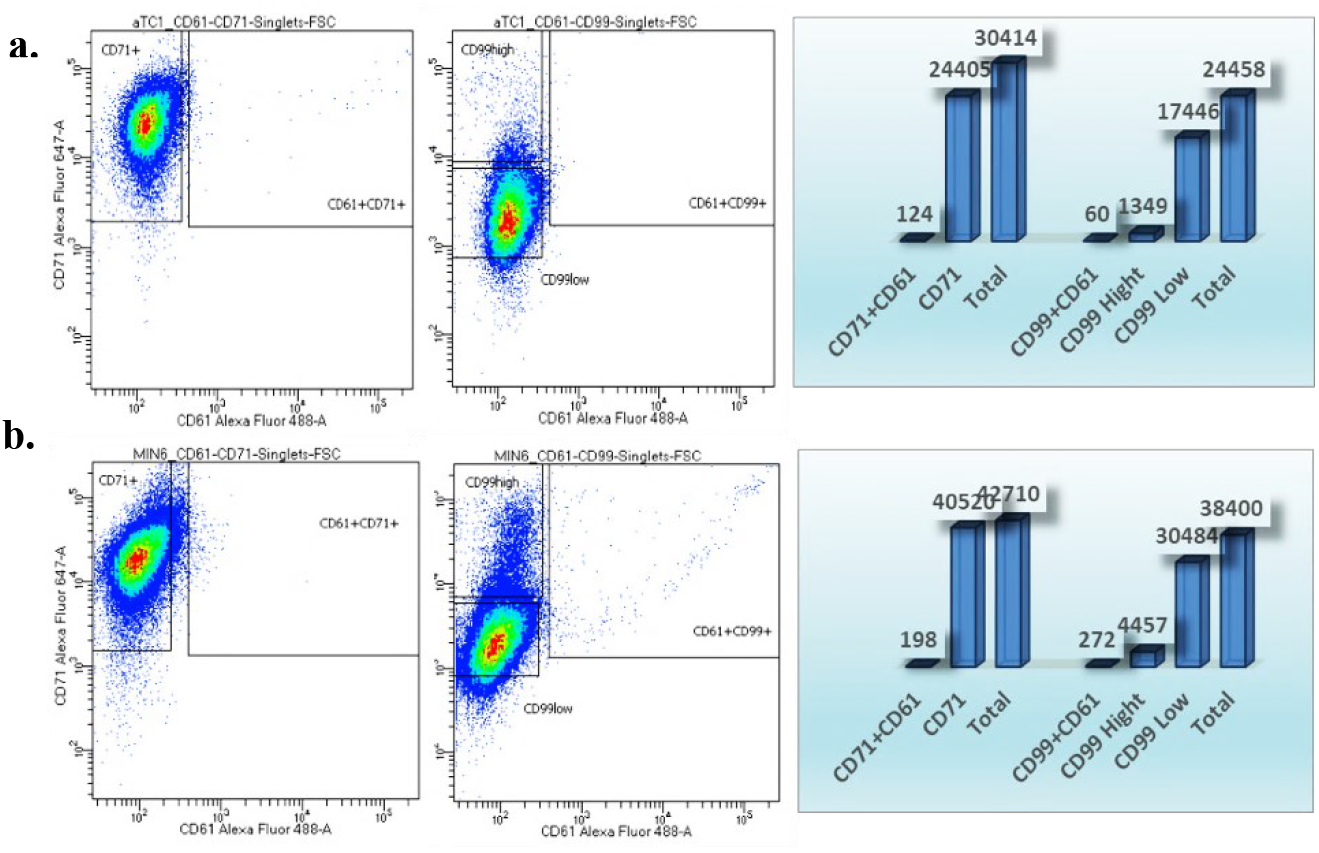
FACS analysis of Cd71, Cd99 and Cd61 expression by MIN6 and αTc1. a) Two subsets are distinguished within αTc1 (Cd99^+^) population: one with high Cd99 expression and a second with low Cd99 expression. In contrast, Cd71 expression seems homogeneous. b) A similar pattern is observed in MIN6 population: Cd71 has homogeneous expression, whereas Cd99 expression varies with one subpopulation with high expression and a second with low Cd99 expression.

**Figure 3:**
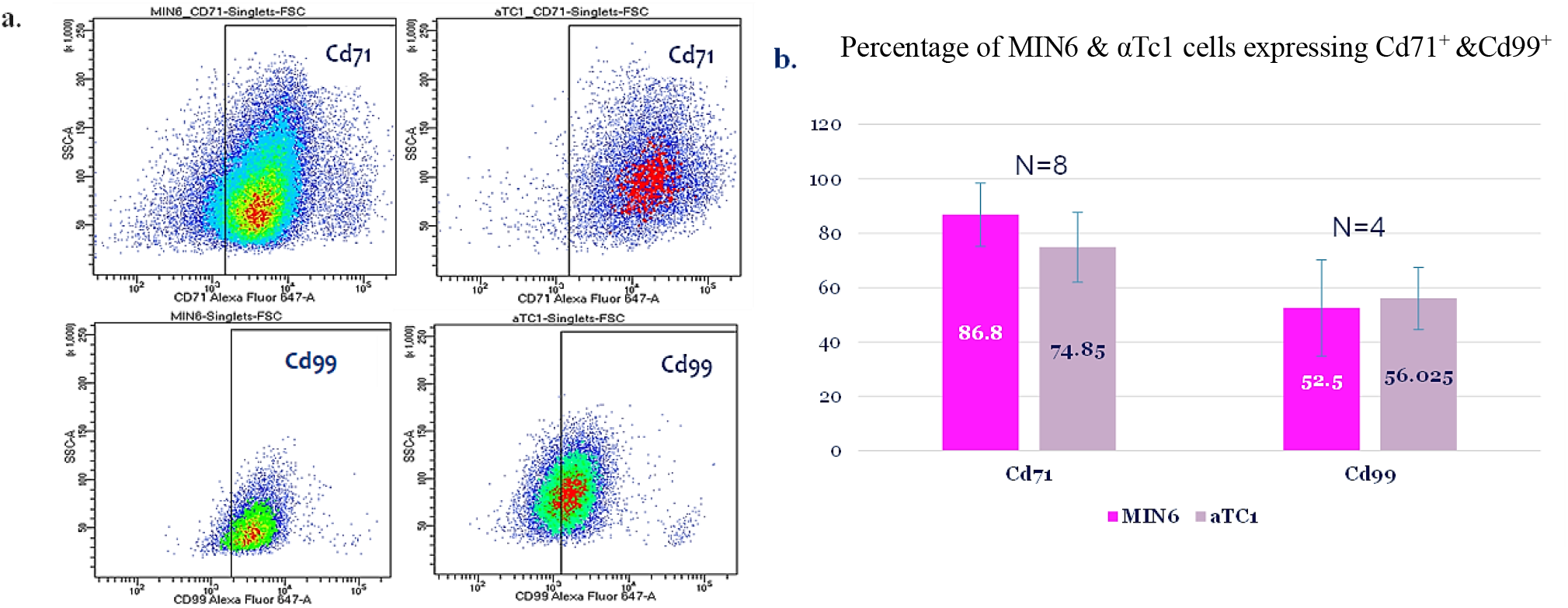
FACS sorting results of MIN6 and α Tc1 expressing Cd71 and Cd99. a) MIN6 and αTc1 cells were labeled with primary anti-Cd71 and anti-Cd99 antibodies (rabbit), then with secondary Alexa-Fluor 647 antibody (anti-rabbit). b) Two subpopulations of MIN6 and αTc1 cells were identified expressing Cd71 and Cd99 receptors. The majority of sorted MIN6 (86,8%) and αTc1 (74.85%) expressed Cd71 compared to 52% of MIN6 and 56% of αTc1 expressed Cd99.

To visualize the proteomic profile of the 144 cells, two principal component analysis were produced, highlighting the existence of proteomic differences between MIN6 and αTc1 cells expressing the same surface epitope. In the PCA (a) four distinct subpopulations are observed when comparing protein expression among the subpopulations of MIN6 (Cd71^+^), MIN6 (Cd99^+^), αTc1 (Cd71^+^), and αTc1(Cd99^+^) (figure 4 a), and in PCA (b) (figure 4 b) by comparing the proteome of the four previously mentioned subpopulations to the MIN6 (P538^+^), MIN6 (Ctl), αTc1 (P538^+^) and αTc1 (Ctl), we can clearly observe eight distinctive subpopulations.

**Figure 4:**
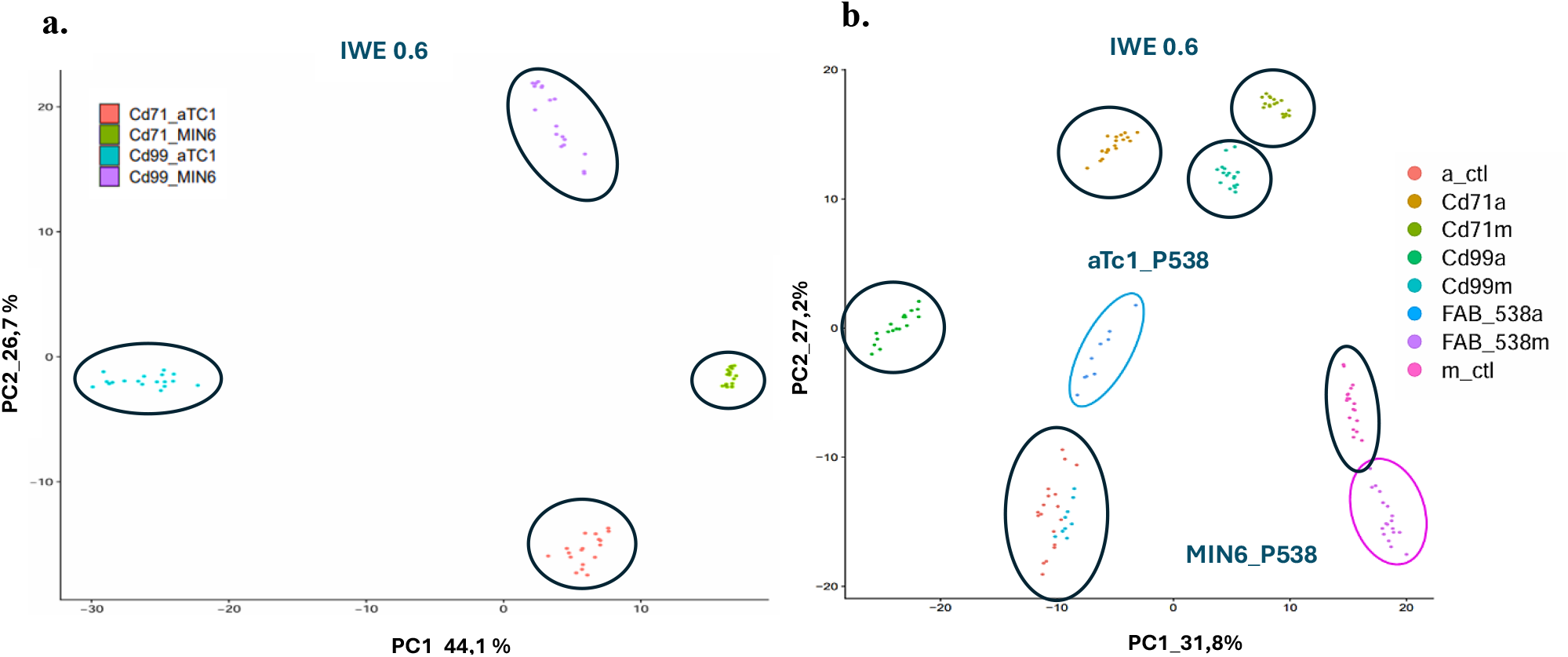
PCA analysis of MIN6 and αTc1 single cell samples. a.) Proteome analysis of 72 cells expressing Cd71 and Cd99 epitopes. Each point represents the proteome of a cell compared to the proteome of other cells. 4 proteomic profiles of cell populations are distinguished by the following colors: Green for MIN6 (Cd71^+^), purple for MIN6 (Cd99^+^), blue for αTc1 (Cd99^+^) and orange for αTc1 (Cd71^+^). b.) Proteome analysis of 144 cells expressing the proteins Cd71, Cd99, P538 and control cells without labeling. Each dot represents the proteome of a single cell compared to the proteome of another cell. The proteomics profile of eight subpopulations is distinguished by their colors: orange for αTc1 (Control), fuchsia for MIN6 (Control), mustard for αTc1 (Cd71^+^), bottle green for MIN6 (Cd71^+^), blue for α Tc1 (P538^+^), purple for MIN6 (P538^+^), light green for αTc1 (Cd99^+^) and light blue for MIN6 (Cd99^+^).

### Distinct expression pattern of the β and α cells hormonal biomarkers

The first group of proteins analysed in the proteomic profile of the eight subpopulations were the pancreatic β and α cells key markers, insulin and glucagon respectively.

#### Glucagon is not an exclusive α cell marker, whereas insulin is specific to β cells

Gcg gene expression was detected in all subpopulations, including MIN6 cells, with higher expression levels in αTc1 subpopulations (figure 5c). In contrast, expression of insulin isoform genes was detected exclusively in the four β cell subpopulations (figure 5a and 5b). The Gcg gene is known for encoding both glucagon and its precursor, proglucagon. In our analysis, the protein code assigned to the Gcg gene by MaxQuant is P55095, which corresponds to proglucagon. According to the work of Perez-Frances et al., Gcg expression is mainly observed in embryonic endocrine cells and their progeny but rarely in adult β-cells (Perez-Frances M, 2022). Indeed, Gcg^+^ embryonic endocrine cells would differentiate predominantly into adult α and δ lineages and would only contribute to about 4% of the adult β cell population (Perez-Frances M, 2022). Eventually, these β cells would continue to express either glucagon or proglucagon even when they reach their adult stage (Perez-Frances M, 2022). On the other hand, Ins^+^ embryonic endocrine cells would exclusively differentiate into adult β cells and no longer be able to express Gcg proteins once mature (Perez-Frances M, 2022). Therefore, the detection of the glucagon precursor, Gcg, in all analyzed MIN6 subpopulations suggests that either the MIN6 lineage are not adult β cells or that all MIN6 subpopulations would come from the same Gcg^+^ embryonic endocrine cells rather than from Ins^+^ embryonic cells.

**Figure 5:**
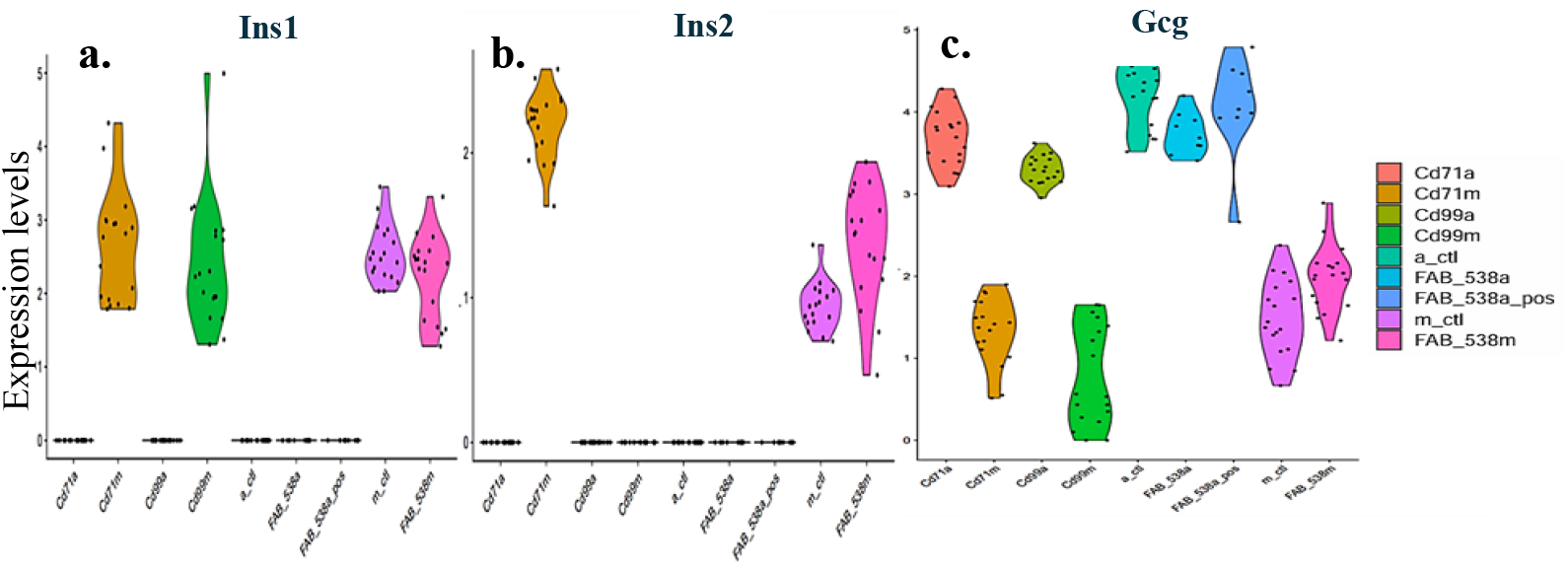
Violon plots of the expression levels of hormonal markers in MIN6 and αTc1 cell populations (CD71^+^, Cd99^+^, P538^+^ and control). a.) Ins1 expression is detected in the MIN6 (Cd71^+^), MIN6 (Cd99^+^), MIN6 (Ctl) and MIN6 (P538^+^) subpopulations. b.) Ins2 expression is detected only in MIN6 (Cd71^+^), MIN6 (Ctl), and MIN6 (P538^+)^ but not in MIN6 (Cd99^+^) subpopulation. c.) Gcg expression is detected in all subpopulations; high levels of Gcg expression are particularly observed in αTc1 (Cd71^+^), αTc1 (Cd99^+^), αTc1 (P538^+^) and α Tc1 (Ctl) cells.

The heterogeneity of MIN6 cells has indeed already been demonstrated in previous work (Zhao R, 2021). However, our results further demonstrate that the MIN6 insulinemic phenotype may originate from Gcg^+^ embryonic endocrine stem cells rather than Ins^+^ stem cells.

#### MIN6 (Cd99^+^) subpopulation lacks the expression of the insulin isoform defining β cells’ functional identity

Compared to glucagon, the insulin hormone in mice has two isoforms, Ins1 and Ins2. All four MIN6 subpopulations expressed Ins1, however, Ins2 was only detected in MIN6 (Cd71^+^), MIN6 (P538^+^), and MIN6 (Ctl), but not in MIN6 (Cd99^+^) single cells and 250 cells of the bulk samples. Studies demonstrated that the deletion of Ins1 has no impact on the function of β cells, whereas Ins2 deletion could result in diabetes development (Skovsø S, 2022). Since the functional identity of β cells is defined by their ability to produce insulin, our findings suggest that the MIN6 (Cd99^+^) subpopulation could have a predisposition to develop a deficiency in insulin production by lacking the expression of Ins2 compared to the other MIN6 subpopulations analyzed, Cd71^+^, P538, and the control group.

### Cells heterogeneity translates through different proteomic profiles across cells expressing the same surface epitopes

Although MIN6 and αTc1 express the same epitopes, functional heterogeneity is observed across the proteome of the eight distinct subpopulations.

#### αTc1(Cd99^+^) cells represent the most homogenous subpopulation

A comparison of the proteomic profiles of MIN6 (Cd99^+^) and αTc1 (Cd99^+^) in the PCA (a) (figure 4 a) revealed significant variations, with the αTc1(Cd99^+^) subpopulation distinguished by the highest number of specific protein groups (49) (table 3 in supplementary materials). This suggests that αTc1 (Cd99^+^) represents the most homogeneous subpopulation compared to the other three, MIN6 (Cd71^+^), MIN6 (Cd99^+^) and αTc1 (Cd71^+^). Furthermore, gene ontology analysis (GO) of the 49 associated proteins revealed that they are primarily involved in metabolic processes, allowing cells to respond to stressful events, like oxidative or genotoxic stress. Some of these biological processes involve diadenosine pentaphosphates, polyphosphates, and inositol phosphates (Ferguson F, 2020). These findings suggest that the αTc1(Cd99^+^) subpopulation compared to MIN6 (Cd99^+^) may have a greater capacity to adapt to stressful environmental conditions.

While 49 protein groups give a homogenous proteomic portrait to the αTc1 (Cd99^+^) subpopulation, the MIN6 (Cd99^+^) subpopulation is characterized by only three proteins: Cck, Sec22b, and Fam120A (figure 7). Of these three proteins, only CcK and Sec22b may play a role in insulin production or secretion (Fan J, 2017) (Cheung GW, 2009).

#### MIN6 (Cd71^+^) proteomic signature is linked to β cells functional identity

when comparing the proteomes of MIN6 and αTc1 cells in the analysis of the principal component a (PCA a), heterogeneity across cell populations expressing Cd71 was equally observed (figure 6). The MIN6 (Cd71^+^) subpopulation is distinguished by the specific expression of Ins2, ERp29, Iapp, Npy, and Tubb2a-Tubb2b. These proteins all play important roles in insulin biosynthesis and are therefore closely associated to the functional identity of MIN6 (Cd71^+^) cells.

**Figure 6:**
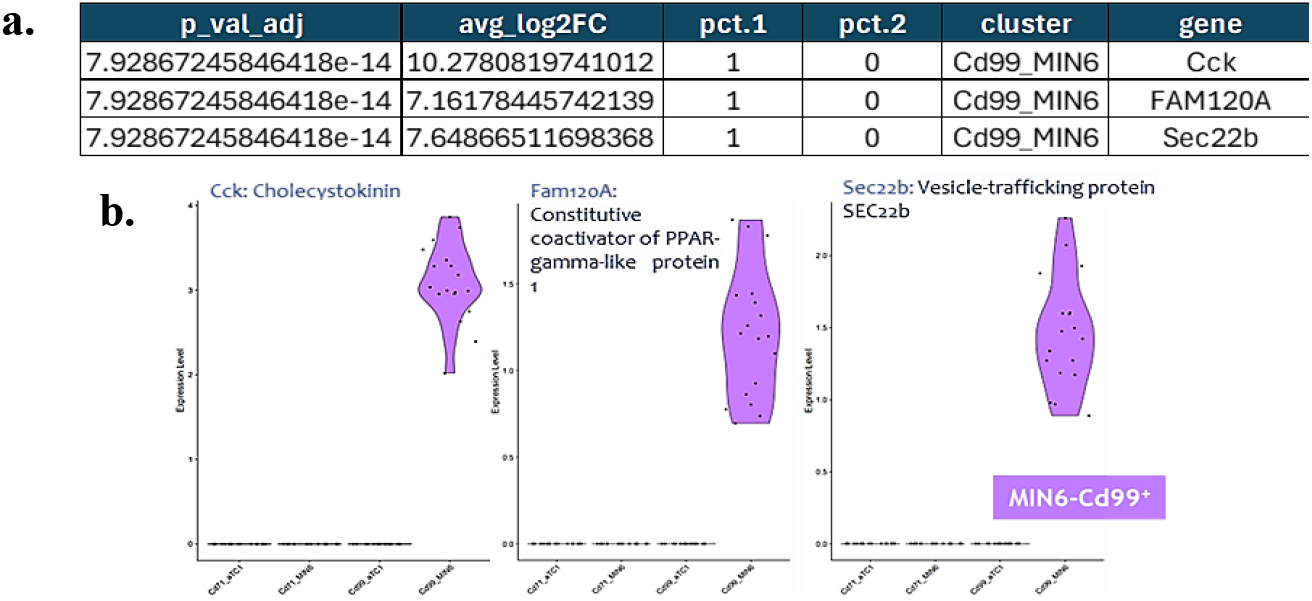
MIN6 (Cd99+) subpopulation-specific protein groups were identified based on Pct.1 = 100%, representing the proportion of cells expressing the protein within the cluster of interest. a.) Table indicating the p values < 0.05 and the avg_Log2FC of each protein, with a Pct.1 = 1 (expression only in MIN6 (Cd99^+^) and a pct. 2 = 0 (no expression in the other subpopulations). b.) Violin plots depicting proteins specifically enriched in the MIN6 (Cd99^+^) subpopulation: Cck, Sec22b and Fam120A.

**Figure 7:**
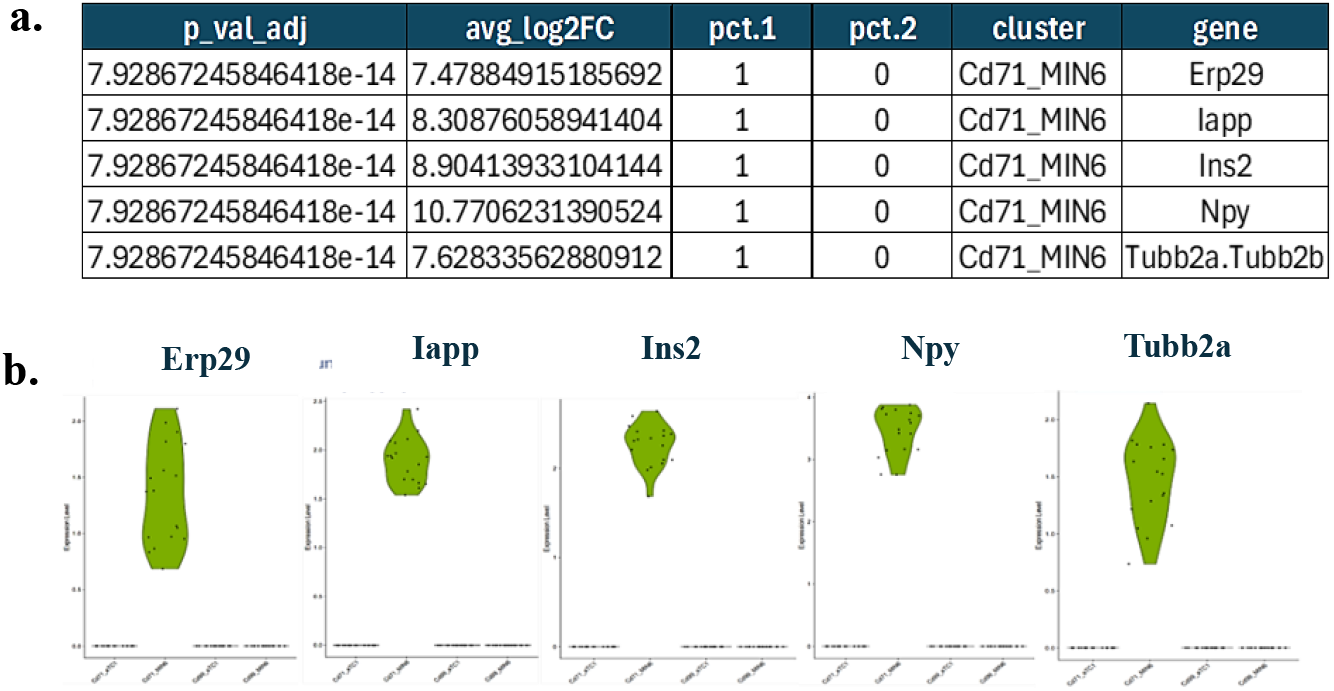
MIN6 (Cd71^+^) subpopulation specific protein groups were identified based on Pct.1 = 100%, representing the proportion of cells expressing the protein within the cluster of interest. a.) Table indicating the p values < 0.05 and the avg_Log2 fold changes of each protein, with a pct.1 = 100% (expression in the MIN6 (Cd71^+^) subpopulation) and a pct. 2 = 0 (no expression in other subpopulations. b.) Violin plots depicting proteins specifically enriched in the MIN6 (Cd71^+^) subpopulation: ERp29, Iapp, Ins2, Npy and Tubb 2a.

In fact, ERp29 is involved in the conversion of proinsulin into insulin (Viviano J, 2020), Iapp is co-secreted with insulin, and its oligomerization has been associated with the development of T2DM (Rehn F, 2024), 2024), Npy acts as a peptidergic neurotransmitter and influences β cells function (Moltz JH, 1985); while Tubb2a and Tubb2b proteins are possible new biomarkers identified in the development of insulin resistance (Pujar MK, 2019).

#### αTC1 (Cd71^+^) subpopulation may play a role in diabetes development in context of iron homeostasis perturbations

While the proteomic profile characterizing MIN6 (Cd71^+^) subpopulation participate in the functional identity of the β cells, the one characterizing the αTc1 (Cd71^+^) subpopulation demonstrates no association to glucagon production or secretion. In fact, the two proteins characterizing αTc1 (Cd71^+^) subpopulation are Sncb and Uqcrp. Sncb (β-Synuclein) is one of the three small acidic proteins of the Synuclein group (α, β, and γ) found to be highly expressed in the brain particularly in the context of Parkinson’s disease and brain tumors (Lavedan C, 1998) (Xiao P, 2023). In pancreatic adenocarcinoma, α-Synuclein has been found to be highly expressed (Geng X, 2010). In the Geng X et al. study, α-Synuclein was also shown to influence insulin secretion by interacting with KATP channels and downregulating insulin granules exocytosis (Geng X, 2010). However, little is known about β-Synuclein expression and function. In our data, however, its expression was detected in αTc1(Cd71^+^), (Cd99^+^) and P538 but not in MIN6 cells subpopulations.

Interestingly, the protein Fth1, which participates in iron storage in the form of ferritin (Hoelzgen F, 2024), was detected exclusively in the αTc1(Cd71^+^) (figure 9). Clinical studies have reported elevated plasma glucagon levels in patients with hemochromatosis (Barton JC, 2017). Given that individuals with hemochromatosis have a high risk of becoming insulin resistant and developing T2D (Barton JC, 2017), our findings suggest that α cells may play a more important role than previously thought in the pathogenesis of T2D, notably in conditions involving iron homeostasis disruption.

**Figure 8:**
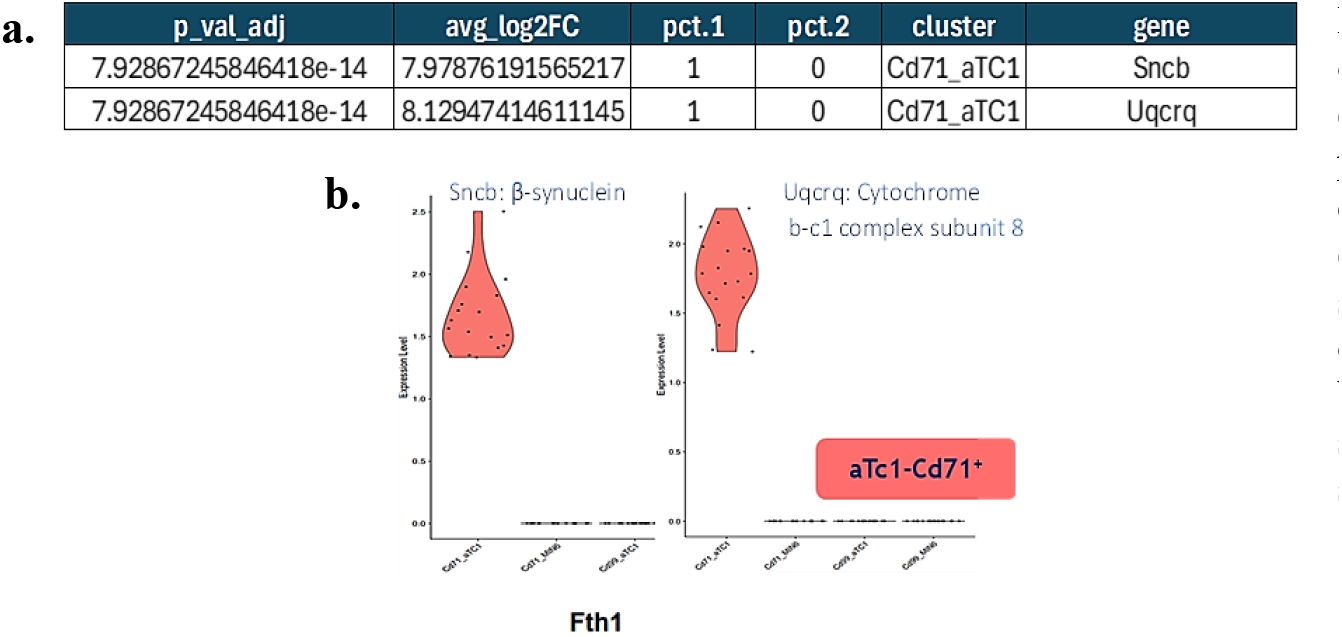
Protein groups expressed only in αTc1 (Cd71^+^) subpopulation (Pct.1 = 100%). **a.)** Table indicating the p values < 0.05 and the aveg_Log2 fold changes of each protein, with a pct.1 = 1 (expression in the αTc1 (Cd71^+^) subpopulation) and a pct. 2 = 0 (absence of expression in other populations). b.) Violin plots depicting proteins specifically enriched in αTc1 (Cd71^+^) subpopulation: Sncb and Uqcrp.

**Figure 9:**
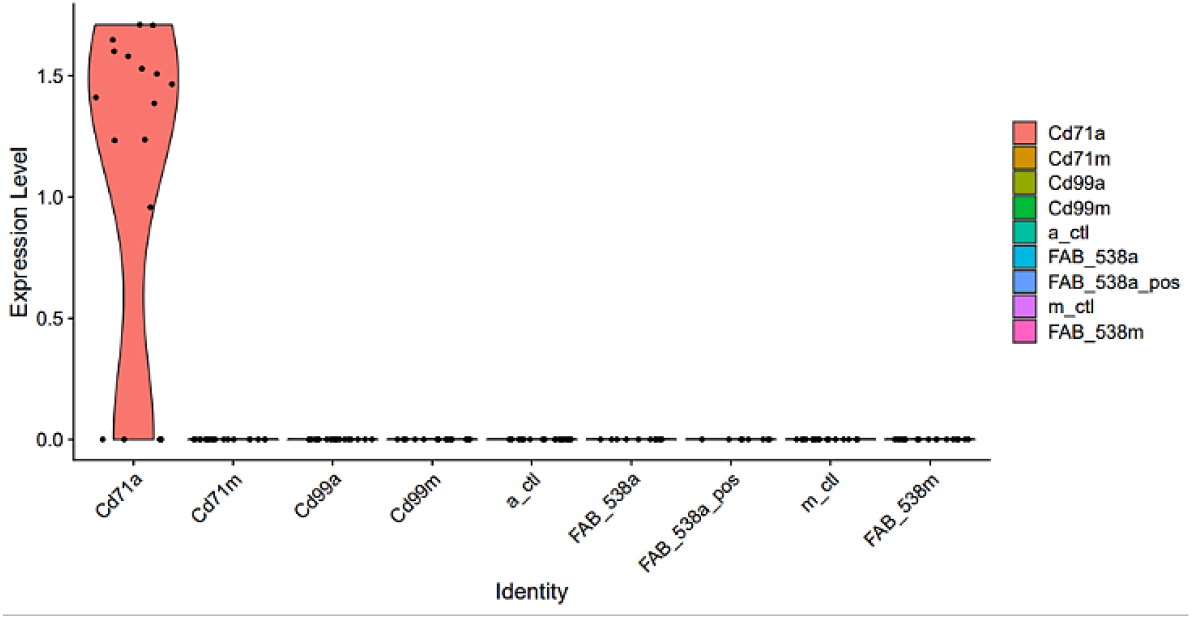
Violin plot of the protein Fth1 (Ferritin heavy chain 1). Fth1 protein expression is not detected in any subpopulation except in αTc1 (Cd71^+^) subpopulation.

### The MIN6 (P538^+^) subpopulation exhibits marked functional-secreting identity compared to other MIN6 and αTc1 subpopulations

β and α cells secrete insulin and glucagon, respectively, which are key hormones characterizing the functional identity of these cell types (Stanojevic V, 2015). The cell membrane proteome of β and α cells is composed of proteins that are part of the secretome (Stutzer I, 2012). The secretome corresponds to a set of proteins secreted by cells through two principal pathways: the exosomal or non-conventional pathway and the classical secretory pathway (Makridakis M, 2010) (Pinheiro-Machado E, 2020). Ongoing research works aim to identify secretome proteins embedded in or linked to plasma membrane (Stutzer I, 2012). However, several proteins involved in vesicular trafficking have already been identified, and a group of these proteins was detected in our proteomic profiling of the MIN6 (P538^+^) subpopulation.

Insulin secretion is a process that takes place in two phases through the classical pathway using granules which are secretory vesicles (Barg S, 2003). The first and short phase, last less than 10 min and allows the insulin secretion from a approximately 1000 vesicles (Barg S, 2003), already docked and primed at the plasma membrane (Yanagisawa Y, 2023) (Barg S, 2003). During this phase insulin is immediately released in response to an increase in glucose (Yanagisawa Y, 2023). The second phase begins immediately after the first and lasts as long as the β cell glucose transporters (GLUTs) are exposed to constant glucose stimulation (Yanagisawa Y, 2023). The Rab-GTPases superfamily is composed of the Rab protein group, in which certain isoforms such as Rab3a and Rab27a play a key role in vesicular trafficking as well as in endo- and exocytosis pathways (Wu SY, 2023) (Veluthakal R, 2023). Rab3a implication is thought to be in the facilitation of the targeting, docking, priming, and fusion of insulin granules with the plasma membrane during exocytosis (Zerial M, 2001) (Pfeffer SR, 2013) (Merrins MJ, 2008) (Norris N, 2024.), and Rab27a has been identified as an important protein in the recruitment and fusion of insulin granules to the membrane (Merrins MJ, 2008).

In our previous study, using bioinformatics prediction analyses with InterProScan, AlphaFold, and AF2Complex, we identified Rab3d (RAB3D, member of the RAS oncogene family) and Ap1m2 (adaptor related protein complex 1 subunit mu 2) as potential candidate proteins for the surface epitope P538, identified using Fab-phage-538 (Moursli Y, 2025). Both Rab3d and Ap1m are part of the protein complexes or families that play essential roles in vesicular trafficking (Collins BM, 2002) (Veluthakal R, 2023) (Wu SY, 2023). Even if the specific role of Rab3d in insulin vesicle trafficking has not been clearly described, this protein belongs to the highly homologous subfamily of Rab3, which includes the 3a, 3b, and 3c isoforms, and their implication in exocytosis regulation has been reported in some studies (Jahn R, 1999). While the Rab-GTPases large family has a ubiquitous expression across all cell types (Pavlos NJ, 2005), the Rab3 subfamily is more specific to secretory cells, such as pancreatic β and α cells (Pavlos NJ, 2005).

Furthermore, when MIN6 cells were incubated in vitro for 48 h in glucose concentrations equivalent to normoglycemia (5.5 mM), Fab-phage-538 binding to MIN6 increased (Moursli Y, 2025). These findings suggest that the expression of P538 increases under conditions where Gluts receptor stimulation is reduced and probably decreases under conditions similar to hyperglycemia (Moursli Y, 2025). Comparable findings were reported for other membrane-associated proteins implicated in vesicle trafficking, such as secretion vesicle v-SNAREs (synaptotagmin and VAMP-2) and target membrane t-SNAREs (syntaxin-1A and SNAP-25), whose expression was decreased in the β cells of animal models affected by T2D (Ostenson H, 2006). Interestingly, the proteome of the MIN6 (P538^+^) subpopulation was distinguished from the other subpopulations by the marked and specific expression of 17 protein groups (figure 10), 12 of which play an important role in vesicular trafficking, exocytosis, insulin secretion, or interact with members of the Rab-GTPases family (figure 11). In the following paragraphs we summarize the potential implication of these 12 protein groups, which are Cd63, DnaJa1, Fkbp11, Glul, Ppp1r1a, Pdxdc1, Nans, Rab8a, Rab6 (a and b), Sec62, Serpinb6, and Scrn1, in insulin production, secretion or even in its action (figure 11).

**Figure 10:**
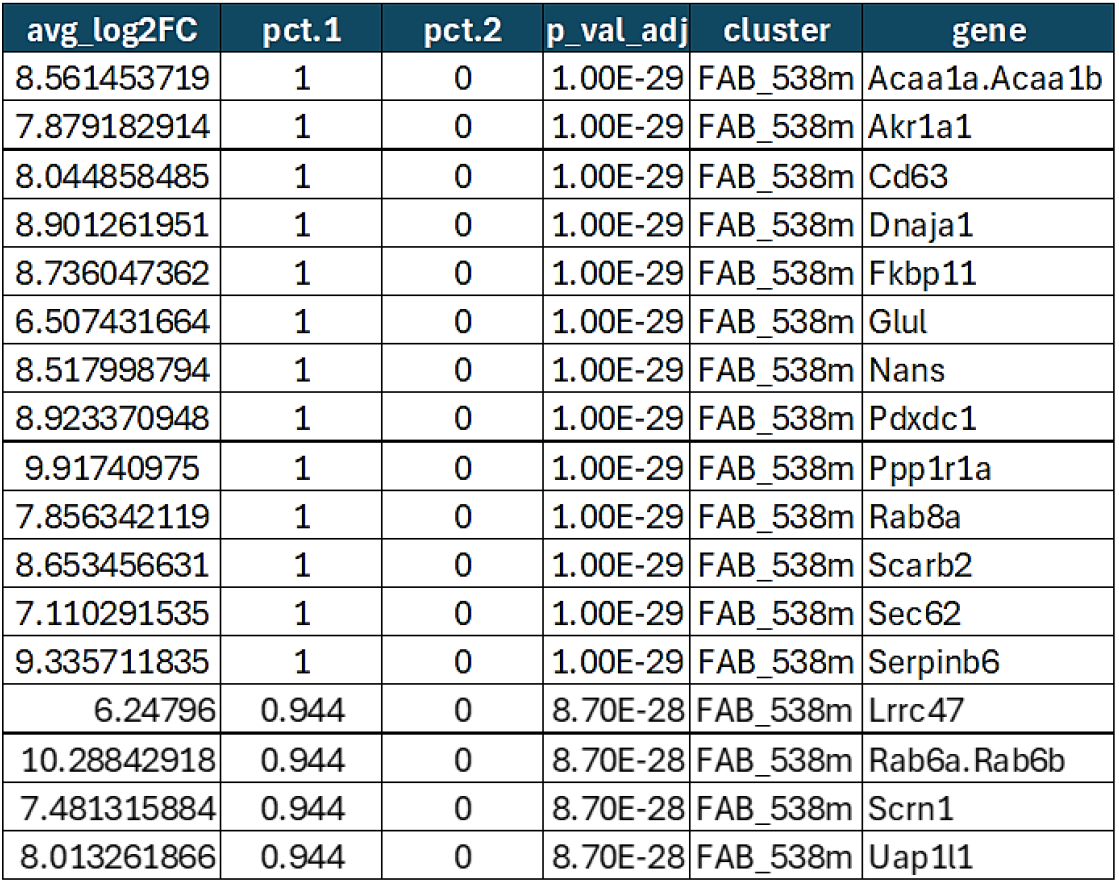
Protein groups expressed only in MIN6 (P538^+^) subpopulation. 17 protein groups with the p values < 0.05 and the aveg_Log2 fold changes of each protein, with a pct.1 ⩾95% (expression in the MIN6 (P538+) subpopulation) and a pct. 2 = 0 (absence of expression in other MIN6 subpopulations of MIN6 and αTc1 expressing Cd71, Cd99, P538 and control)

**Figure 11:**
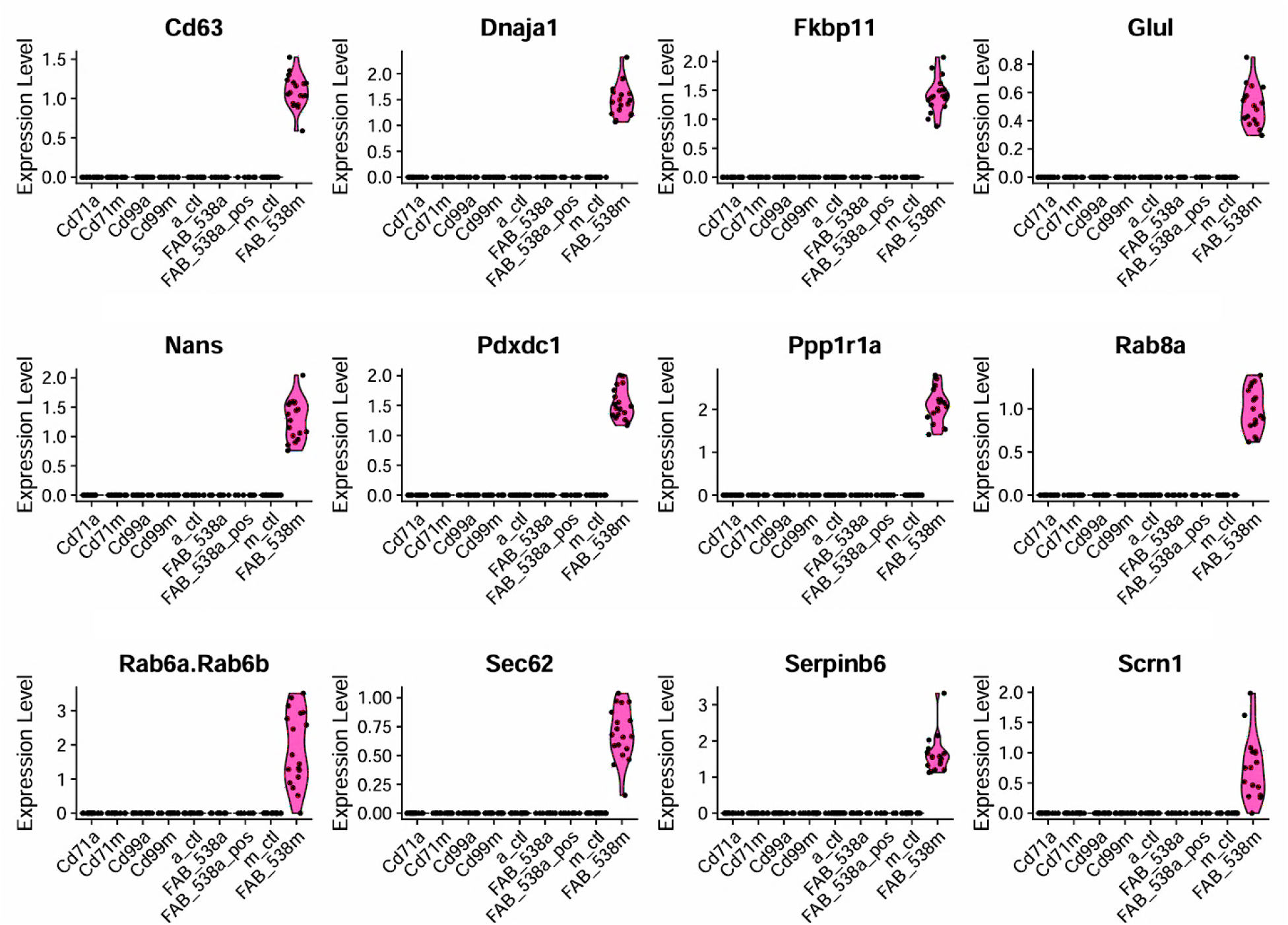
Violin plots of the 12 proteins specific to the MIN6 (P538^+^) subpopulation and directly or indirectly involved in vesicular trafficking, insulin production or secretion. The high expression of these 12 proteins has been observed exclusively in MIN6 (P538^+^) subpopulation (FAB_538m) in pink color of the Violin plots.

Cd63 is a tetraspanin protein that interacts with Rab-GTPase family, particularly with Rab isoforms Rab5c, Rab7a, Rab10, Rab11, and Rab31 (Cheerathodi M, 2021). Furthermore, both Cd63 expression and insulin secretion were reported to increase in human and murine pancreatic β cells under glucose stimulation, suggesting that Cd63 is implicated in insulin secretion (Rubio-Navarro A, 2023). Conversely, the same team reported that in the pathological condition of diabetes, there is a decrease in β cells expressing Cd63, as β cells lacking Cd63 expression are unable to secrete insulin (Rubio-Navarro A, 2023).

DnaJa1 is part of the J-domain proteins (JDPs) group, and it facilitates the binding and the activation of Hsp70 ATPase through the J domain (Braun JEA, 2023). DnaJa1 was detected in small extracellular vesicles of neuronal tissues, suggesting its involvement in vesicular trafficking (Braun JEA, 2023) (Brenna S, 2020). Although DnaJa1 detection at the level of insulin vesicles has not yet been demonstrated, the DnaJ group is involved in insulin secretory granule maturation (GSI) (Ungewickell E, 1995) (Omar-Hmeadi M, 2021). In fact, the process of proinsulin conversion into insulin requires the dissociation of the clathrin covering the GSI (Omar-Hmeadi M, 2021). This step involves heat shock proteins, notably Hsp70 and auxilin, which is part of the DnaJ group or Hsp40 family (Kirchhausen T, 2014) (Ungewickell E, 1995) (Gruschus JM, 2004).

Furthermore, glutamine synthetase (Glul) is an enzyme participating in insulin secretion by converting glutamine into glutamate, which strengthens the calcium signaling needed to amplify insulin secretion induced by glucose stimulation (Han G, 2021). Ppp1r1a is a phosphatase involved in Langerhans islet function regulation (Taneera J., 2015) (Taneera J, 2024). An increase in Ppp1r1a expression has been observed in association with an elevation of insulin secretion suggesting that Ppp1r1a can be considered as a biomarker of β cell dysfunction in both humans and mice (Taneera J., 2015) (Taneera J, 2024). Pdxdc1 is a decarboxylase enzyme (Pyridoxal dependent decarboxylase domain containing 1), and its human equivalent, PDXDC, is attached to the Golgi apparatus and vesicles in several tissues including pancreatic cancer cells, the CAPAN-2 (The Human Protein Atlas., 2025), but not specifically in β cells. The data we have collected indicate that this protein is highly expressed specifically in the MIN6 (P538^+^) subpopulation, with an average Log2FC of 8.9 (figure 10).

The Nans (*N*-acetylneuraminate synthase) is an enzyme that, according to the Human Protein Atlas, its human equivalent, NANS, is mainly localized in the nucleoplasm and the plasma membrane (The Human Protein Atlas., 2025). It plays a crucial role in vesicular trafficking through the synthesis of sialic acid proteins, namely the Neu5Ac and Neu5Gc (Kavaler S, 2011). Neu5Ac is used by the Golgi apparatus, during the process of glycoconjugation, once activated into CMP-Neu5Ac (Tran C, 2021). Kavaler et *al*. showed that a deficiency in Neu5Ac in mice resulted in the development of glucose intolerance and a fasting hyperglycemia due to pancreatic β cells disfunction confirmed by 40% decrease in insulin secretion *in vivo* (Kavaler S, 2011).

Rab8a, Rab6a, and Rab6b, all members of the Rab-GTPase group, are important regulators of vesicular trafficking in all secretory cells, including pancreatic β cells (Grigoriev I, 2007). By binding to the cytoplasmic and the Golgi apparatus vesicles, they guide vesicular insulin transport between the Golgi apparatus, the ER, and the plasma membrane (Grigoriev I, 2007) (Fye MA, 2023). Sec62, a translocation protein and member of the chaperone proteins such as Hsp70 and Calmodulin, is one of the membrane proteins of the ER (Liu M, 2018). Thus, Sec62 plays an important role in the process of insulin maturation, since it facilitates the post-translational translocation of preproinsulin (Liu M, 2018). The Scrn1 implication in vesicle trafficking has essentially been studied at the presynaptic level of neurons (Lindhout FW, 2019). However, its specific role in insulin exocytosis or vesicular trafficking has not yet been clearly demonstrated.

Taken together, the implication of the principal protein groups characterizing the MIN6 (P538^+^) subpopulation and the possible predicted proteins identifying P538, in vesicular trafficking, exocytosis, insulin maturation, and secretion suggests that this newly identified subpopulation may hold a more specialized functional identity compared to the other MIN6 (Cd71^+^), MIN6 (Cd99^+^), and control subpopulations.

Moreover, these results support our initial hypothesis that two cell lines derived from the same tissue and expressing the same surface epitope can exhibit heterogeneity in their proteomic profile. Indeed, while the specific expression of 17 protein groups confers to the MIN6 (P538^+^) subpopulation an insulin production and secretion functional identity, αTc1 (P538^+^), on the other hand, was characterized by only two proteins, Pof1b (Premature ovarian failure 1B) and Pkp1 (Plakophilin-1), with no established involvement in glucagon secretion.

Also, while the MIN6 (Cd99^+^) subpopulation may be predisposed to the development of insulin deficiency compared to MIN6 (Cd71^+^) and MIN6 (P538^+^), the proteomic profile of αTc1 (Cd99^+^) subpopulation suggests that this subpopulation may hold a greater ability to adapt to stressful environmental conditions. Finally, when comparing MIN6 (Cd71^+^) to αTc1 (Cd71^+^), our findings suggest that the MIN6 subpopulation has probably a lesser role in T2DM genesis in the context of iron homeostasis disruption than αTc1 (Cd71^+^).

## Limitations of the study

Single cell proteomic methods are a fascinating research field and hold great promise for understanding how cell diversity can be leveraged to solve several pathological mechanisms. However, these approaches are still subject to some limitations that may have an impact on both the number and type of proteins detected in one cell and therefore influence data analysis and proteomic profile interpretation. These variations are usually explained by different factors, and many of which are technical, particularly in our study.

### Cell line types

The number of protein groups that can be detected and characterized in a single cell is not an absolute standard number, fixed and identical to all cell types, and therefore may vary depending on the origin of the tissue and animal species (Petelski AA, 2021).

### The total number of single cells analyzed

It is well known that a larger number of single cells increases the statistical power of a study (Slavov N, 2022). In fact, this allows not only to identify variations between cell populations as discrete as they might be but also to enhance the accuracy of principal component analysis (PCA) (Slavov N, 2022) (Adil A, 2021).

### Material loss during sample preparation

Several single cell methods face the risk of proteomic material loss during sample processing. In this regard, automated systems and miniaturization allow to control and reduce the variability between samples and losses (Petelski AA, 2021) (Petelski AA., 2022). Indeed, given that the concentrations of proteomic material of a single cell, as well as that obtained after trypsinization, are very low, the risk of adsorption loss on surfaces is all the higher when the wells receiving the cells are large and the sorting volumes are significant (Kelly RT, 2020) (Wu R, 2019).

### The data acquisition mode

The mode of data acquisition can affect in a non-predictable way the performance of analytical algorithms used in the analysis of proteomic data (Slavov N, 2020) (Wallmann G, 2023). The SCoPE2 method is based on data independent acquisition (DIA) (Wallmann G, 2023), whereas the method used in this study was data dependent acquisition (DDA). In DDA, the instrument’s peptide detection parameters are programmed to selectively fragment the most abundant precursor ions (Ghosh G, 2025) (Guo J, 2020). This could be a source of bias regarding the groups of proteins identified and analyzed, because the instrument analyzes only the most abundant peptides by omitting to analyze those whose abundance falls below a threshold for detecting precursor ions conducive to fragmentation (Ghosh G, 2025) (Guo J, 2020). In contrast, in DIA there is a simultaneous fragmentation of all precursor ions whose mass is within a defined mass interval (Wallmann G, 2023) (Guo J, 2020) (Ghosh G, 2025). Therefore, it is expected that all protein groups identified will be most representative of the actual proteome expressed by the cell, without excluding groups with low abundance. This advantage is one of the arguments that justified the use of DIA parameters by several research groups, including the Slavov team (Wallmann G, 2023) (Slavov N., 2021).

### Effect of carrier channel with isotopic labeling like TMT

Using carriers in each mixture of single cells certainly allows better identification of peptides, but it can also mask the signal of weakly abundant peptides because carrier peptide signals are much stronger than that of single cells (Goto-Silva L, 2021). This could bias the quantification or even generate background noise, thus limiting the identification of peptides and additional proteins (Goto-Silva L, 2021).

## Conclusion

Diabetes is one of the oldest health conditions described in medical history, with its incidence and prevalence continuing to rise globally. Several complications affecting the cardiovascular, neurological, and renal systems are associated with it. In the last decade, the heterogeneity of pancreatic β cells in the context of diabetes has been reported in some studies. Single cell transcriptomic and genomic analysis have considerably deepened our understanding of this cell diversity. However, to better understand the differences in cells functional identity and the variations that characterize them, the study of the proteome is necessary. Single-cell proteomic methods allow establishing a correlation between cells’ proteomic profiles and the expression of membrane proteins, thereby distinguishing cell populations based on their functional attributes. This level of precision is highly relevant to therapeutically target specific dysfunctional cell subpopulations. Precision medicine is the future, and the use of more specific and targeted treatments will undoubtedly require a better mastery and understanding of the particularities that define the diversity of cell populations not only through their genomes and transcriptomes but also through their changing proteomic profiles at the single cell level.

## Supporting information

Supplementary materiel

## Acknowledgement

We thank other members of the Coulombe laboratory for helpful discussions. We acknowledge the use of OpenAI’s ChatGPT (GPT-4) for language refinement and grammar corrections.

