## Supplementary materiel for "Discovery and characterization of a pancreatic β cell subpopulation expressing an unknown surface epitope through single cell proteomics"

| **MaxQuant parameters (v2.6.5.0)** | **Sections** |
| --- | --- |
| 1. Loading raw data | Raw data |
| 2. Identification of the experiments | Raw data |
| 3. Reporter MS2 and select the type of TMT plex (10-plex/11-plex) | Group specific settings   - Group 0: type |
| 4. Edit the correction factors for the type of TMT plex |  |
| 5. Selection of isotopic weight = IWE 0.6 | Group specific settings   - Groupe 0: Misc |
| 6. Trypsine | Group specific settings   - Groupe 0: digestion mode |
| 7. Fixed modification: remove carbamidomethylation (C). | Group specific settings   - Groupe 0: digestion mode |
| 8. Add: select Fasta (Swissprot-Mousse file), identification rules 9. Table:   • MS scan table  • Table m/z | Global settings:   - Sequences |
| 10. PSM-FDR = 0.01 11. Proteins-FDR = 0.01 | Identification |
| 12. Select matches between experiments (select match between runs) |  |
| 13. Save |  |

**Table 1**: Parameters used in the reporter ion intensity analysis with MaxQuant (v2.6.5.0).

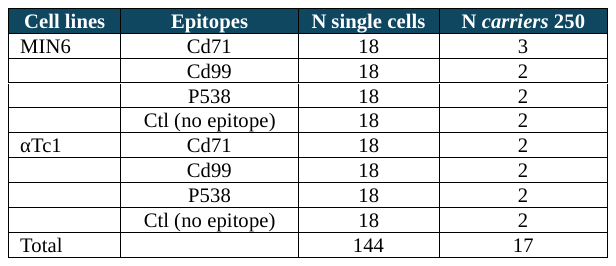

**Table 2:** Number of single cell proteomic experiments performed with samples MIN6 and αTc1 expressing (Cd71^+^), (Cd99++), P538^+^ and Ctl.

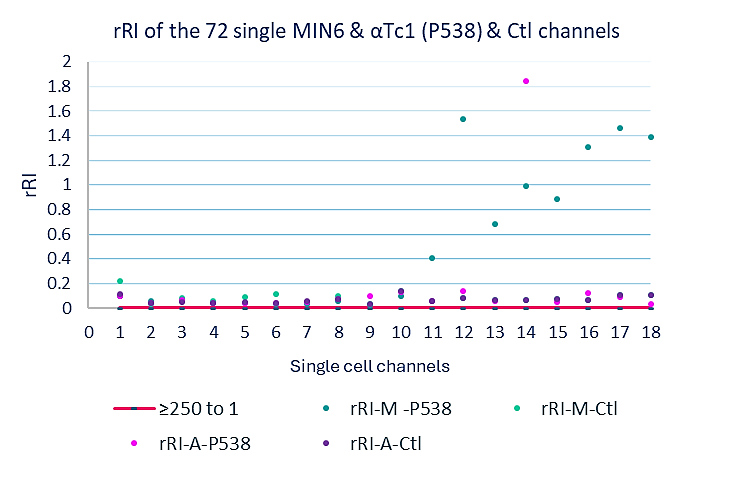

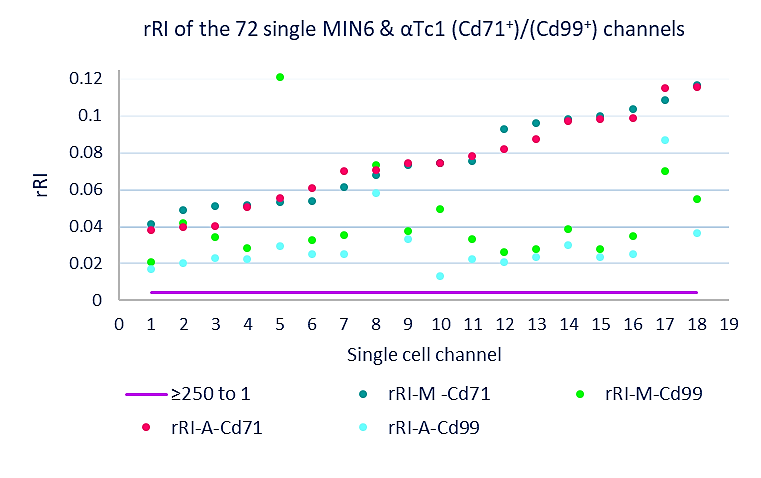

**a.**

**b.**

**Figure 1: relative reporter intensity (rRI) values distribution per TMT channel of the 144 single MIN6 and αTc1 cells expressing Cd71^+^, Cd99^+^, P538^+^, and Ctl .** The rRI of all cells are greater than the expected threshold of 1: 250, or 0.004.

**Table 3: List of 49 proteins groups expressed exclusively in αTc1 (Cd99^+^) (Cd99_aTc1) cell subpopulation:** 49 proteins, whith adjusted p_value < 0.05 indicate statistical significance, are exclusively expressed in the group of αTc1 (Cd99^+^) (pct. 1 = 100%, pct. 2 = 0). The differential analysis of log2 fold change (avg_Log2FC) shows that all these proteins have a marked overexpression with log2 FC values greater than 5.

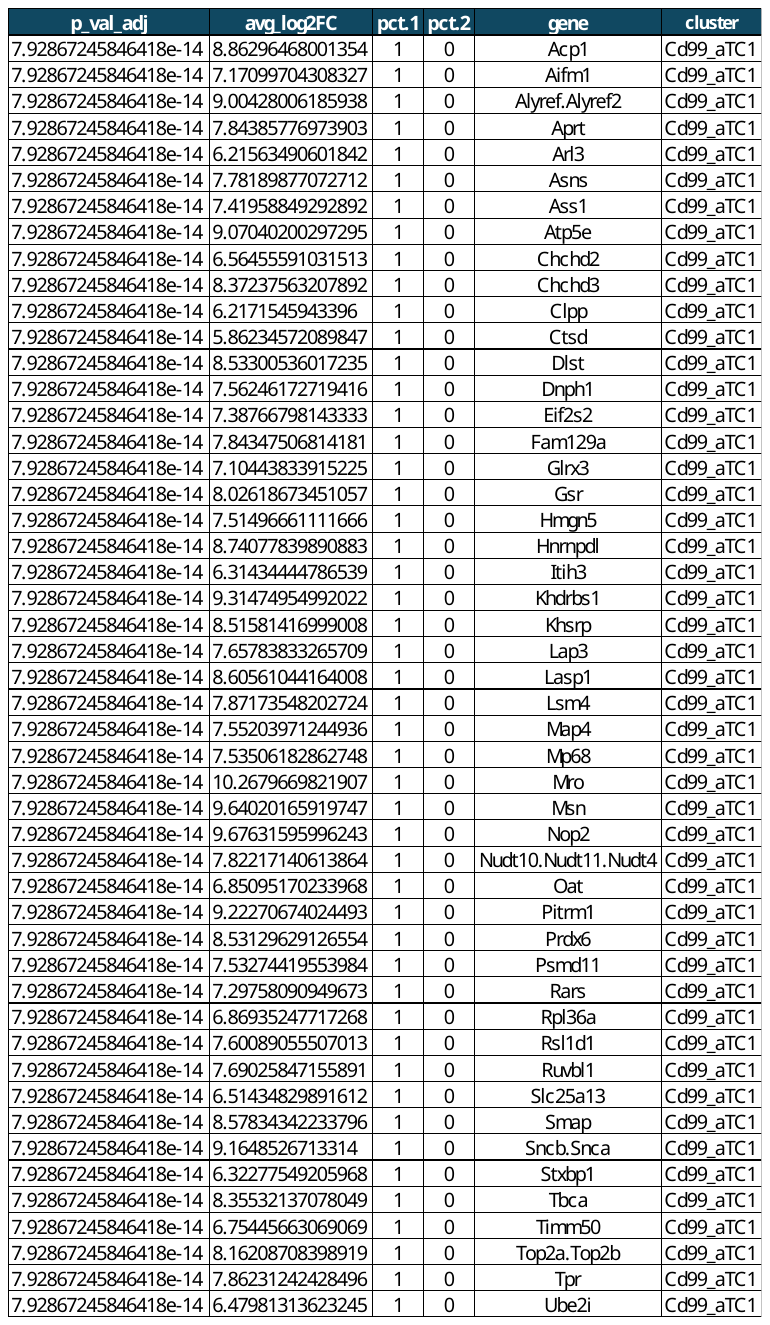
